# Predicting Endometriosis Status and Menstrual Cycle Phase Using DNA Methylation

**DOI:** 10.64898/2026.09.02.749016

**Authors:** Amrita Nagasuri, Umair Khan, Parker Grosjean, Adi Siddharth, Idit Kosti, Sally Mortlock, Sahar Houshdaran, Nilufer Rahmioglu, Stacey A. Missmer, Krina T. Zondervan, Grant Montgomery, Christian M. Becker, Peter Rogers, Juan Irwin, Tomiko Oskotsky, Karla Lindquist, Christopher Seaman, Linda C. Giudice, Marina Sirota

**Affiliations:** Bakar Computational Health Sciences Institute, University of California, San Francisco, CA, USA; The Institute for Molecular Bioscience, The University of Queensland, Brisbane, QLD, 4072, Australia; Australian Women and Girls’ Health Research Centre, School of Public Health, The University of Queensland, Brisbane, QLD, 4006, Australia; Department of Obstetrics, Gynecology and Reproductive Sciences, University of California, San Francisco, CA, USA; Department of Epidemiology, Harvard T.H. Chan School of Public Health, Boston, MA, USA; Boston Center for Endometriosis, Boston Children’s Hospital and Brigham and Women’s Hospital, Boston, MA, USA; Division of Adolescent and Young Adult Medicine, Department of Medicine, Boston Children’s Hospital and Harvard Medical School, Boston, MA, USA; Department of Obstetrics, Gynecology, and Reproductive Biology, College of Human Medicine, Michigan State University, Grand Rapids, MI, USA; Centre for Human Genetics, University of Oxford, Oxford, UK; Oxford Endometriosis CaRe Centre, Nuffield Department of Women’s and Reproductive Health, John Radcliffe Hospital, University of Oxford, Oxford, UK; University of Melbourne Department of Obstetrics and Gynaecology, Royal Women’s Hospital, Melbourne, Australia; Department of Pediatrics, University of California, San Francisco, San Francisco, CA, USA; Department of Epidemiology and Biostatistics, University of California, San Francisco, CA, USA

## Abstract

Endometriosis is a chronic inflammatory disease associated with pelvic pain, infertility, and delayed diagnosis. Growing evidence suggests that altered DNA methylation contributes to disease development and could serve as a biomarker for disease. We developed a leakage-safe machine learning pipeline to classify endometriosis case-control status and menstrual cycle phase using genome-wide DNA methylation data from eutopic endometrial tissue. The dataset consisted of 984 samples profiled using the Illumina Infinium MethylationEPIC array, with measurements across approximately 759,000 CpG sites. Technical variation was corrected using SmartSVA batch correction. Ridge logistic regression models were trained using stratified 80/20 train-test splits, with regularization strength selected via stratified cross-validation. Feature selection approaches included ridge coefficient ranking, per-CpG t-tests, and univariate logistic regression with FDR correction. Model validity was evaluated using label-shuffling analyses. Menstrual cycle phase classification showed strong performance (mean cross-validation AUROC: 0.971, held-out test AUROC: 0.989), reflecting genome-wide hormonally driven methylation. Ridge regression produced lower but meaningful performance for endometriosis classification (mean cross-validation AUROC: 0.854, held-out test AUROC: 0.875). Ridge coefficient-based feature selection identified compact predictive CpG sets, supporting the hypothesis that endometriosis-associated methylation signal is distributed across many loci rather than a few highly predictive CpGs. Pathway enrichment analyses identified substantial enrichment for menstrual cycle phase but limited enrichment for disease status following FDR correction, consistent with a diffuse endometriosis-associated signal. These findings demonstrate that ridge regression can detect methylation patterns associated with both endometriosis and menstrual cycle phase, highlighting the importance of accounting for cycle-related epigenetic variation in endometrial DNA methylation studies.

**Graphical Abstract:** 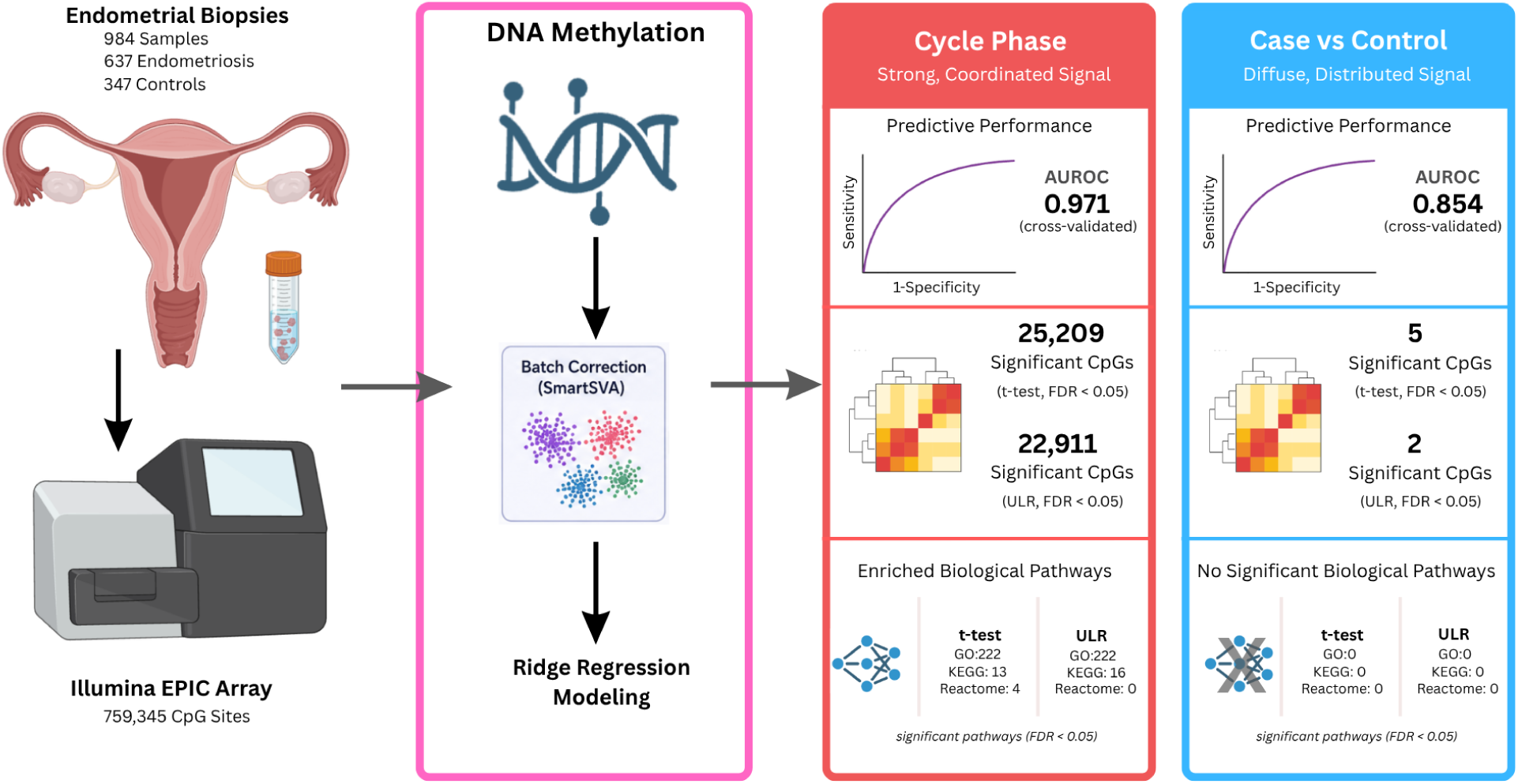

## Introduction

Endometriosis is a chronic inflammatory gynecologic disease characterized by the growth of endometrial-like tissue outside the uterus. It affects approximately 10% of reproductive-age women and is commonly associated with chronic pelvic pain, dysmenorrhea, dyspareunia, infertility, and reduced quality of life (Zondervan et al., 2020). Increasing evidence suggests that endometriosis should be viewed as a complex systemic disease involving inflammation, immune dysregulation, fibrosis, angiogenesis, and altered hormonal signaling rather than solely the presence of ectopic lesions (Saunders and Horne, 2021). Despite its prevalence, diagnosis remains challenging and often requires invasive surgical confirmation, resulting in an average diagnostic delay of approximately seven years from symptom onset (Shafrir et al., 2018). The disease is also highly heterogeneous, with considerable variation in symptom severity, disease stage, lesion characteristics, and reproductive outcomes. This clinical heterogeneity has complicated efforts to identify consistent molecular signatures and develop reliable diagnostic biomarkers.

Evidence suggests that molecular alterations associated with endometriosis extend beyond ectopic lesions and are also present within the eutopic endometrium. In addition to ectopic disease, the eutopic endometrium of some affected individuals exhibits altered cellular and molecular function, including progesterone resistance and impaired endometrial receptivity, suggesting that it represents a biologically relevant tissue for investigating disease-associated molecular alterations (Burney and Giudice, 2012). Among the mechanisms implicated in disease pathogenesis, epigenetic regulation has emerged as an important area of study. DNA methylation, a modification involving the addition of a methyl group to cytosine residues at CpG dinucleotides, plays a central role in regulating gene expression and cellular function. In the endometrium, methylation patterns are not static and change throughout the menstrual cycle in response to hormonal signaling and tissue remodeling. As a result, the menstrual cycle phase represents a major source of biological variation in endometrial methylation studies and can complicate the identification of disease-associated signals. The development of high-throughput platforms such as the Illumina Infinium MethylationEPIC array now allows methylation to be measured across hundreds of thousands of CpG sites, providing an opportunity to examine these relationships at a genome-wide scale.

The most comprehensive characterization of DNA methylation in the eutopic endometrium was recently reported by Mortlock et al., whose study forms the foundation of the present work. Using the Illumina Infinium MethylationEPIC array, the authors analyzed genome-wide DNA methylation across 984 eutopic endometrial samples from women with and without endometriosis. They demonstrated that menstrual cycle phase is the dominant source of endometrial DNA methylation variation, identifying more than 9,600 differentially methylated CpG sites between proliferative and secretory endometrium, with enriched pathways consistent with endometrial remodeling throughout the menstrual cycle. In contrast, methylation differences between endometriosis cases and controls were comparatively modest, with no CpG sites reaching genome-wide significance for overall disease status, although stronger methylation signals emerged in stage III/IV disease. Despite these subtle case-control differences, genome-wide DNA methylation collectively explained approximately 24% of the variance in endometriosis status, with approximately 16% remaining after accounting for genetic effects. Together, these findings suggest that disease-associated information is distributed throughout the endometrial methylome, while also highlighting the limitations of conventional single-CpG analyses for detecting subtle, heterogeneous methylation patterns.

Machine learning approaches have increasingly been applied to endometriosis diagnosis and biomarker discovery as a means of integrating complex clinical and molecular datasets. A recent systematic review by Zhang et al. identified 45 studies applying machine learning across clinical, imaging, genetic, and other omics data, demonstrating the growing use of these approaches for endometriosis diagnosis while also highlighting the need for improved standardization and external validation. Similarly, machine learning has been successfully applied to transcriptomic datasets to identify combined diagnostic biomarkers through feature selection, predictive modeling, and validation in independent cohorts, demonstrating the ability of multivariate approaches to extract predictive molecular signatures from high-dimensional molecular data. Despite these advances, relatively few studies have applied machine learning to genome-wide DNA methylation data from the eutopic endometrium. Given that disease-associated methylation signals appear to be subtle and distributed across many CpG sites, multivariate machine learning approaches may complement conventional epigenome-wide association analyses by leveraging predictive information across the methylome, providing the rationale for the present study.

Analyzing genome-wide methylation data presents challenges due to the large number of measured CpG sites relative to the number of available samples. Biologically relevant information may be distributed across many CpGs rather than driven by a small number of strongly associated sites. Machine learning approaches may help identify these complex patterns and may capture signals that are difficult to detect using traditional univariate methods alone. In this study, we investigated methylation patterns associated with both menstrual cycle phase and endometriosis status using a large cohort of eutopic endometrial samples (Figure 1A). We evaluated predictive performance using regularized regression models, compared multiple feature selection strategies, performed pathway enrichment analyses, and examined the extent to which multivariate and univariate approaches capture methylation variation.

**Figure 1.**
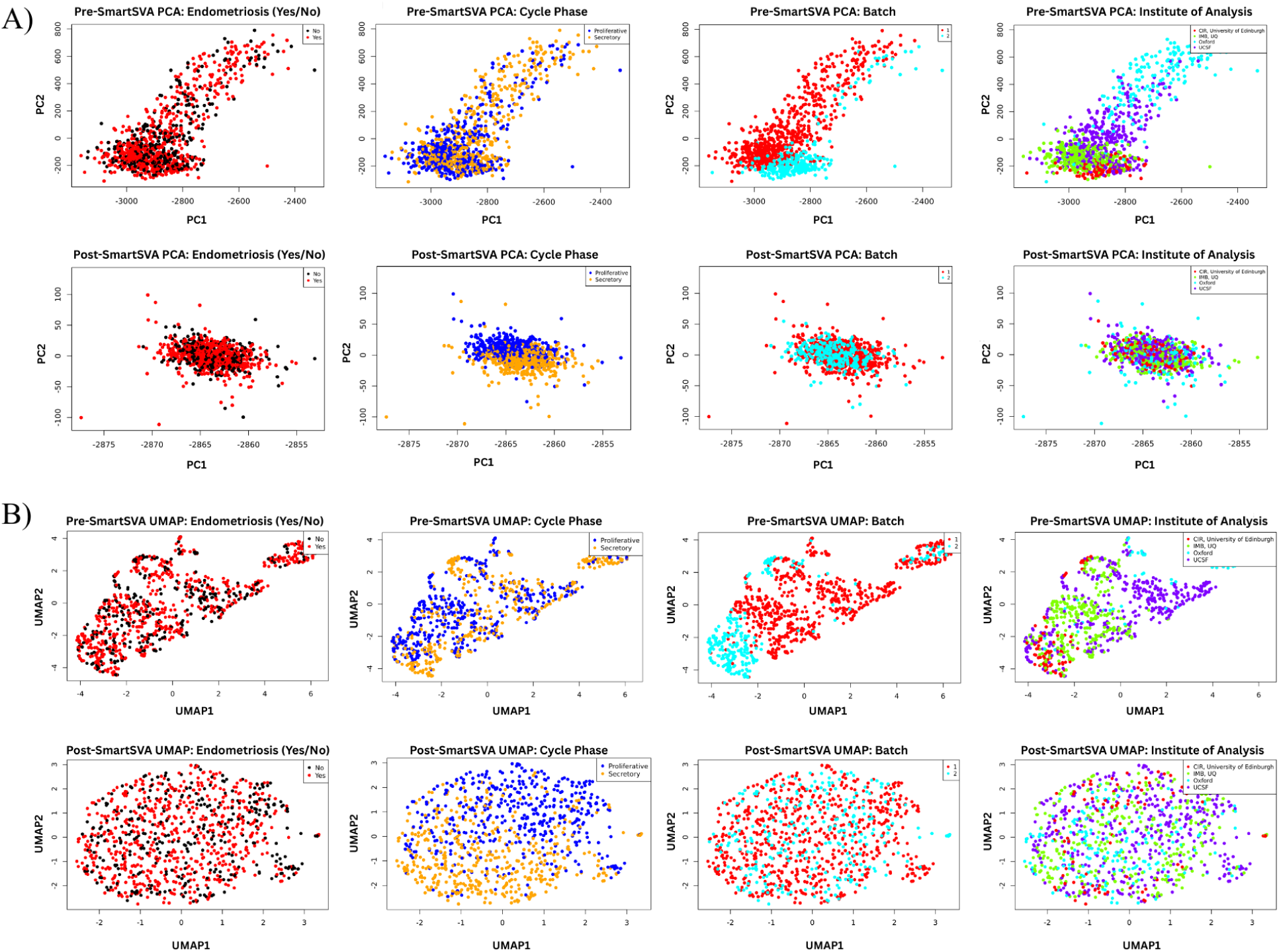
A) Overview of the study. B) PCA and C) UMAP visualizations before and after SmartSVA correction demonstrating reduction of technical clustering while preserving biological variation. Analyses were performed using 984 eutopic endometrial samples profiled across 759,345 CpG sites following quality control. Respective PCA and UMAP plots are colored by endometriosis status, menstrual cycle phase, institute of analysis, and batch before and after SmartSVA correction. SmartSVA substantially reduced clustering associated with batch and institute of analysis while preserving biological variation associated with menstrual cycle phase and, to a lesser extent, endometriosis status.

## Methods

### Data

We analyzed a dataset consisting of 984 eutopic endometrial biopsy samples profiled across multiple independent studies. The cohort included 637 patients with surgically and histologically confirmed endometriosis and 347 controls without endometriosis, including 201 controls classified as having no uterine or pelvic pathology (NUPP). Cases were further annotated by disease stage, including stage I/II, stage III/IV, and a small number of samples with unknown stage classification. Menstrual cycle phase was assigned in the original study using the histologic criteria of Noyes et al., as described by Mortlock et al. (2023). Because proliferative substages were not available across all contributing cohorts, proliferative samples were consolidated into a single proliferative endometrium (PE) category. Samples were annotated by menstrual cycle phase as proliferative endometrium (PE), early secretory (ESE), mid-secretory (MSE), late secretory (LSE), secretory endometrium without further subclassification (SE), and menstrual. For binary cycle phase classification analyses, PE samples were assigned to the proliferative group, whereas ESE, MSE, LSE, and SE samples were combined into a single secretory group. Menstrual samples were excluded from cycle phase classification analyses.

Available metadata included age, BMI, parity, age at menarche, genetic ancestry, disease stage, lesion type, pain phenotypes, batch, and institute of analysis. Genome-wide DNA methylation data were generated using the Illumina Infinium MethylationEPIC BeadChip array. Quality control and preprocessing were performed by Mortlock et al. using their published pipeline, including probe- and sample-level filtering to remove low-quality measurements before downstream analysis (Mortlock et al., 2023). We used the resulting quality-controlled dataset as the starting point for our analyses. Menstrual-phase samples were excluded only from binary menstrual cycle phase analyses. Methylation levels were represented using M-values, calculated as the logit transformation of beta values, and were used for all downstream analyses. The final methylation matrix consisted of 759,345 CpG sites measured across 984 eutopic endometrial samples. Samples originated from studies conducted at the University of Oxford, the University of Melbourne, the University of Edinburgh, and the University of California, San Francisco. Clinical metadata were standardized across cohorts, and endometriosis case-control labels and menstrual cycle phase annotations were consolidated into consistent categories.

### Filtering Criteria

For endometriosis case-control analyses, all 984 samples passing quality control were retained. For menstrual cycle phase classification, the prediction task was restricted to binary classification of proliferative vs secretory cycle phases. Samples annotated as menstrual phase (n = 50) were excluded because they could not be assigned to either the proliferative or secretory class. Endometriosis cases with unknown disease stage (n = 7) were also included, resulting in a final dataset of 934 samples consisting of 473 proliferative and 461 secretory endometrial samples.

### Batch Correction

The quality-controlled methylation dataset was first divided into stratified training (80%) and held-out testing (20%) subsets. SmartSVA correction was then performed separately for the training and test datasets to prevent information leakage between the two subsets. Surrogate variables for the training data were estimated using a model that included the outcome of interest, whereas correction of the held-out test data was performed using a null model that did not include test-set outcome labels. The corrected training data were used for feature selection, hyperparameter tuning, and model fitting, while the independently corrected test data were reserved for final model evaluation. Dataset structure before and after SmartSVA correction was evaluated using silhouette scores, with principal component analysis and UMAP used to visualize changes in clustering. Visualizations were colored by batch, institute of analysis, endometriosis status, and menstrual cycle phase to assess the reduction of technical variation while preserving biological signal. Silhouette scores were used to quantify changes in clustering associated with technical and biological variables. Although applying SmartSVA separately to the training and test subsets may result in slightly different latent technical factors being estimated in each subset, this strategy was chosen to prevent information from the held-out test samples from influencing model development.

### Predictive Modeling

Following the stratified train-test split and separate SmartSVA correction of the training and test subsets, model development was performed exclusively within the training data. The held-out test set remained isolated from feature selection, model training, and hyperparameter tuning and was used only for final model evaluation. Ridge logistic regression models with L2 regularization were selected because previous work suggested that endometriosis-associated methylation differences are distributed across many correlated CpG sites rather than concentrated within a small number of highly informative loci. Regularization strength was optimized by evaluating 25 logarithmically spaced values of the inverse regularization parameter (C), ranging from 10^-6^ to 10^2^. The optimal parameter was selected using stratified cross-validation within the training data. Feature-selection procedures were performed using the training data, with per-CpG t-tests and univariate logistic regression explicitly restricted to the training subset prior to evaluation of reduced models. Model performance was evaluated using AUROC and AUPRC during cross-validation and by independent prediction on the held-out test set using the complete methylation dataset of 759,345 CpG sites.

To assess whether model performance exceeded chance, outcome labels were randomly shuffled 10 times for each classification task. For each shuffle, samples were divided into stratified 80/20 training and test sets, with five-fold stratified cross-validation performed within the training set and performance evaluated on the held-out test set. Shuffled-label analyses used the full CpG feature set without coefficient-based feature selection. For case-control classification, the regularization parameter was selected across the same 25-value C grid for each shuffle, whereas cycle-phase classification used the regularization parameter from the primary model.

### Feature Selection

Multiple feature-selection strategies were used to identify informative CpGs associated with each classification task. Supervised feature selection ranked CpGs according to the absolute magnitude of coefficients estimated from ridge logistic regression models fitted using the training data. The ranked coefficient distributions were evaluated using KneeLocator, and the detected elbow was used to define reduced feature sets. Independent two-sided Welch’s t-tests and univariate logistic regression analyses were performed across all 759,345 CpG sites using the training data. Statistical significance was determined using Benjamini-Hochberg false discovery rate correction, with an adjusted P value below 0.05 considered statistically significant. Heatmaps were generated using CpG sets identified through ridge coefficient ranking, t-tests, and univariate logistic regression. Methylation values were standardized using row-wise z-score normalization, and hierarchical clustering was performed using Euclidean distance and complete linkage. Sample annotations indicated endometriosis status and menstrual cycle phase.

### Pathway Analysis

Pathway enrichment analysis assessed the biological relevance of selected CpG feature sets. CpGs were mapped to genes and analyzed using the missMethyl package through the gometh framework using Gene Ontology (GO), Kyoto Encyclopedia of Genes and Genomes (KEGG), and Reactome pathway databases. Statistical significance was evaluated using Benjamini-Hochberg FDR correction. Analyses were performed in R (version 4.5.2) using missMethyl (version 1.42.0).

### Statistical Analyses

Statistical analyses were performed using the quality-controlled methylation dataset described above. Endometriosis case-control analyses included all 984 samples, while menstrual cycle phase analyses included 934 samples after exclusion of 50 menstrual-phase samples.

Predictive models were developed using stratified 80/20 training and held-out test splits, with five-fold stratified cross-validation performed within the training data. Model performance was evaluated using AUROC and AUPRC. Per-CpG analyses were performed using two-sided Welch’s t-tests and univariate logistic regression, with statistical significance determined using Benjamini-Hochberg false discovery rate correction at an adjusted P value below 0.05. To evaluate whether predictive performance exceeded chance, outcome labels were randomly shuffled 10 times for each classification task and model performance was evaluated using the shuffled labels as described above.

### Code availability

Code used for data preprocessing, statistical analyses, predictive modeling, feature selection, pathway analysis, and visualization is publicly available in the Endometrial Methylation Prediction GitHub repository: https://github.com/anagasuri/Endometrial-Methylation-Prediction.

### Use of large language models

OpenAI ChatGPT was used during manuscript preparation to assist with organization, restructuring, and refinement of written content and with debugging of analysis code. All scientific interpretations, analytical decisions, results, and conclusions were independently reviewed and verified by the authors. ChatGPT was not used to generate data or independently conduct the analyses reported in this study.

## Results

The final dataset included 984 eutopic endometrial samples, consisting of 637 endometriosis cases and 347 controls (Table 1). Among the cases, 344 samples were classified as stage I/II disease, 286 as stage III/IV disease, and 7 had unknown stage information. Samples represented all annotated menstrual cycle phases (PE, ESE, MSE, LSE, SE, and menstrual) and originated from multiple contributing cohorts. For binary cycle phase analyses, PE samples were classified as proliferative, ESE/MSE/LSE/SE samples were grouped as secretory, and menstrual samples were excluded. PCA and UMAP analyses demonstrated substantial reduction of clustering associated with batch and institute following SmartSVA correction (Figure 1B and C; Supplementary Figure S1). Silhouette score analysis supported these observations, showing decreased clustering by technical variables after correction (Supplementary Table S1). In contrast, variation associated with menstrual cycle phase remained evident, indicating that cycle-phase-related structure was preserved. Endometriosis-associated variation was less apparent after correction.

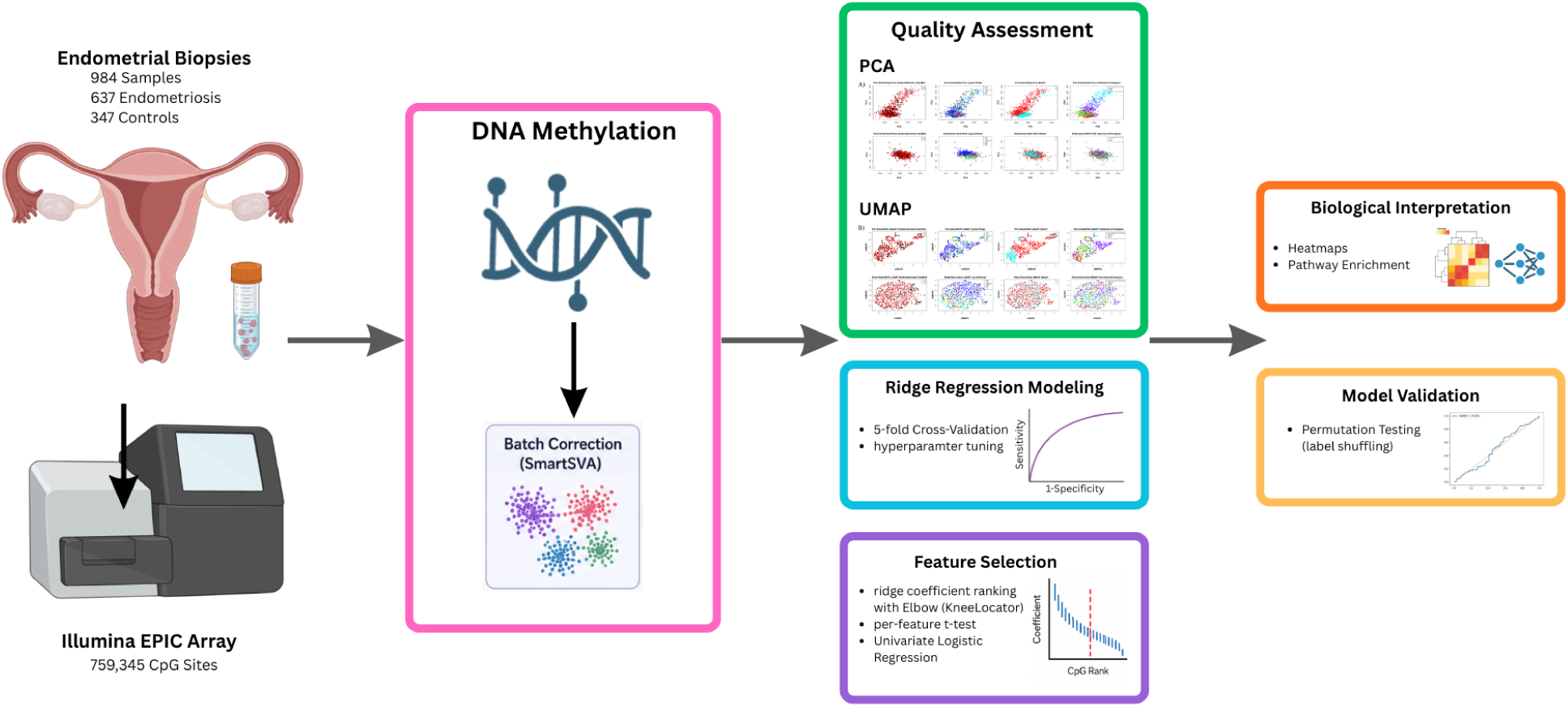

**Table 1.** Characteristics of the study cohort used for DNA methylation analyses. A total of 984 eutopic endometrial biopsy samples (637 endometriosis cases and 347 controls) passed quality control and were included in downstream analyses. Samples are summarized by endometriosis status, menstrual cycle phase, disease stage, institute of analysis, and processing batch. Values are presented as n (%).

| Characteristic | Overall | Controls | Endometriosis |
| --- | --- | --- | --- |
| Sample size | 984 | 347 | 637 |
| Menstrual cycle phase (original annotation) |  |  |  |
| PE | 473 (48.1%) | 170 (49.0%) | 303 (47.6%) |
| ESE | 122 (12.4%) | 44 (12.7%) | 78 (12.2%) |
| MSE | 209 (21.2%) | 78 (22.5%) | 131 (20.6%) |
| LSE | 108 (11.0%) | 35 (10.1%) | 73 (11.5%) |
| SE | 22 (2.2%) | 7 (2.0%) | 15 (2.4%) |
| Menstrual | 50 (5.1%) | 13 (3.7%) | 37 (5.8%) |
| Menstrual cycle phase (binary analysis) |  |  |  |
| Proliferative | 473 (48.1%) | 170 (49.0%) | 303 (47.6%) |
| Secretory | 461 (46.8%) | 164 (47.3%) | 297 (46.6%) |
| Excluded (menstrual) | 50 (5.1%) | 13 (3.7%) | 37 (5.8%) |
| Endometriosis stage (cases only) |  |  |  |
| Stage I/II | 344 (54.0%) | — | 344 (54.0%) |
| Stage III/IV | 286 (44.9%) | — | 286 (44.9%) |
| Unknown | 7 (1.1%) | — | 7 (1.1%) |
| Institute for analysis |  |  |  |
| CIR, University of Edinburgh | 83 (8.4%) | 31 (8.9%) | 52 (8.2%) |
| IMB, UQ | 312 (31.7%) | 87 (25.1%) | 225 (35.3%) |
| Oxford | 178 (18.1%) | 52 (15.0%) | 126 (19.8%) |
| UCSF | 411 (41.8%) | 177 (51.0%) | 234 (36.7%) |
| Batch |  |  |  |
| 1 | 707 (71.8%) | 260 (74.9%) | 447 (70.2%) |
| 2 | 277 (28.2%) | 87 (25.1%) | 190 (29.8%) |

### Menstrual Cycle Phase Is Associated with Robust, Genome-Wide DNA Methylation Remodeling and Near-Perfect Classification Performance

Ridge regression trained on the full methylation dataset (759,345 CpGs) classified proliferative and secretory phase samples with high accuracy. After excluding menstrual samples (n = 50), the binary cycle phase analysis included 934 samples consisting of 473 proliferative and 461 secretory endometrial samples.

The full-feature ridge logistic regression model achieved a mean cross-validation AUROC of 0.971 and a mean cross-validation AUPRC of 0.965 within the training data. When evaluated on the independently held-out test set, the model achieved an AUROC of 0.989 and an AUPRC of 0.990 (Figure 2A). Classification performance substantially exceeded that observed for endometriosis status and remained consistent across validation folds. Shuffled-label analyses produced AUROC values near 0.5, indicating that the observed predictive performance was not reproduced after randomization of the outcome labels. These findings were consistent with the PCA, UMAP, and silhouette score analyses, which identified menstrual cycle phase as the strongest source of biological variation following batch correction. Together, these results indicate that methylation patterns associated with cycle phase consistently distinguish proliferative and secretory endometrium.

**Figure 2.**
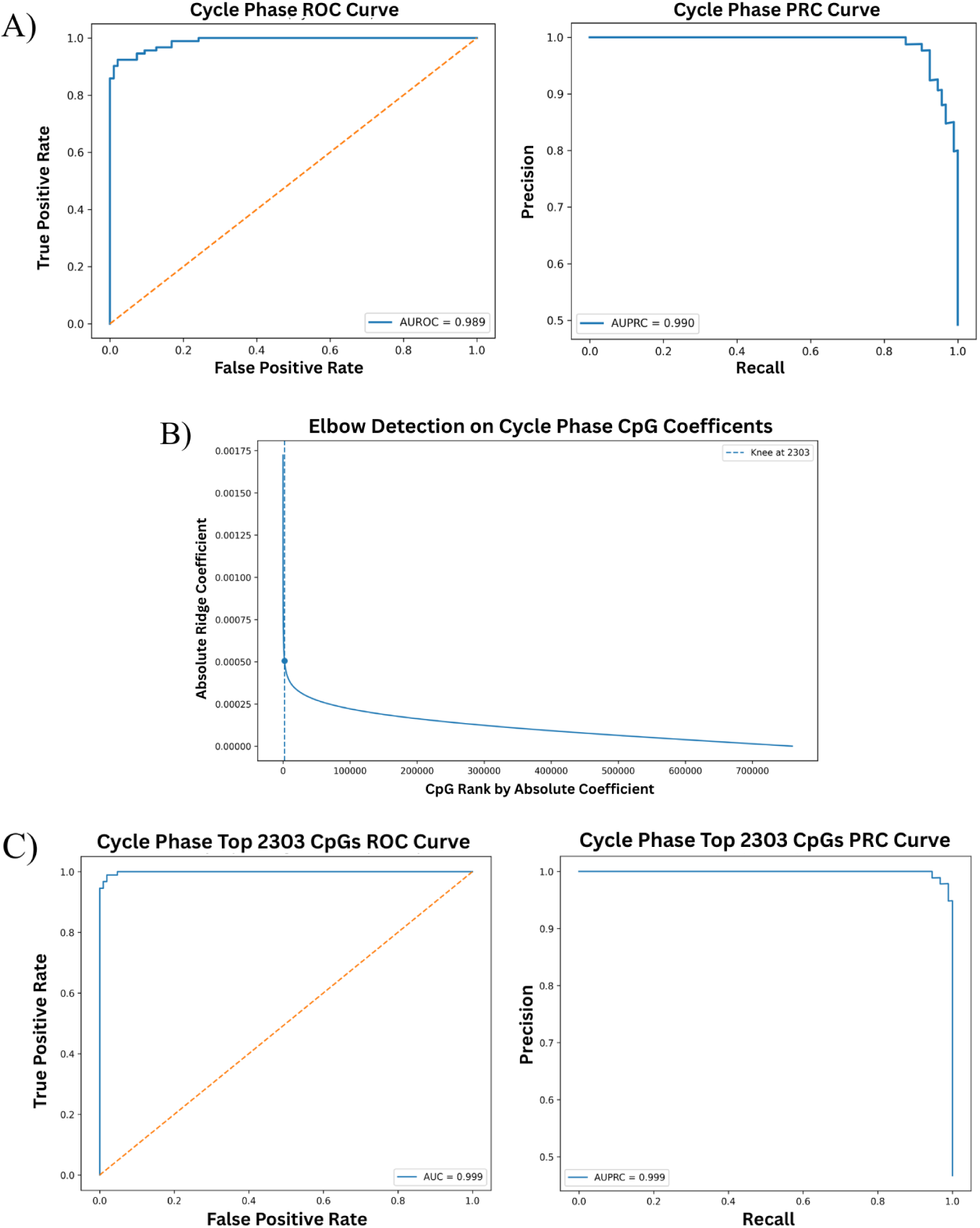
Predictive performance of cycle phase using ridge logistic regression models using the full genome-wide DNA methylation feature set. A) ROC and PRC curves are shown for classification of menstrual cycle phase using all available CpG sites following quality control and SmartSVA correction. The cycle phase model achieved a held-out test AUROC of 0.989 and a held-out test AUPRC of 0.990, indicating near-perfect separation of proliferative and secretory phase samples. These results demonstrate that the menstrual cycle phase represents a substantially stronger source of methylation variation in eutopic endometrial tissue than endometriosis status. B) Elbow point identification for ridge regression coefficient–based feature selection. Absolute ridge regression coefficients were ranked from largest to smallest for the menstrual cycle phase model using the full genome-wide methylation feature set. The elbow point was identified using the KneeLocator algorithm to identify where ridge coefficients began to decrease more gradually. The detected elbow corresponded to 2,303 CpG sites for the cycle phase model, indicated by the dashed vertical line. This CpG subset was selected as a reduced feature set for downstream modeling and biological interpretation. C) Performance of reduced-feature ridge regression models using CpG subsets selected by coefficient-based elbow analysis. ROC and PRC curves are shown for the ridge logistic regression model trained using the top 2,303 CpG sites identified from the menstrual cycle phase model. The reduced-feature cycle phase model achieved a held-out test AUROC of 0.999 and a held-out test AUPRC of 0.999, demonstrating that nearly all predictive information was retained after reducing the feature set from 759,345 to 2,303 CpGs.

Ranking CpGs by absolute ridge coefficient magnitude revealed a gradual decline in feature importance rather than a sharp transition point. Elbow detection identified a reduced feature set consisting of 2,303 CpGs (Figure 2B). Independent univariate analyses produced substantially larger feature sets, with 25,209 CpGs passing FDR significance thresholds in t-test analyses and 22,911 CpGs passing false discovery rate significance thresholds in univariate logistic regression analyses. Although the number of significant CpGs differed across feature selection methods, all approaches identified large numbers of cycle phase-associated methylation sites. These results indicate that proliferative and secretory endometrium differ across thousands of genomic loci rather than a limited set of isolated CpGs.

A ridge logistic regression model trained using the 2,303 ridge-selected CpGs retained predictive performance comparable to that of the full-feature model. The reduced-feature model achieved a mean cross-validation AUROC of 0.971 and a mean cross-validation AUPRC of 0.965 within the training data. When evaluated on the independently held-out test set, the model achieved a held-out test AUROC of 0.999 and a held-out test AUPRC of 0.999 (Figure 2C). These findings indicate that cycle phase classification performance was retained after reducing the feature space from 759,345 CpGs to 2,303 CpGs.

Heatmaps generated from ridge-selected, t-test-selected, and univariate logistic regression-selected CpGs all demonstrated clear separation between proliferative and secretory phase samples (Figure 3). Samples clustered primarily according to cycle phase regardless of feature selection method. Similar clustering patterns were observed across all three heatmap approaches despite large differences in feature count. The consistency of these patterns indicates that cycle phase-associated methylation differences occur across many CpG sites rather than being restricted to a small number of highly influential loci.

**Figure 3.**
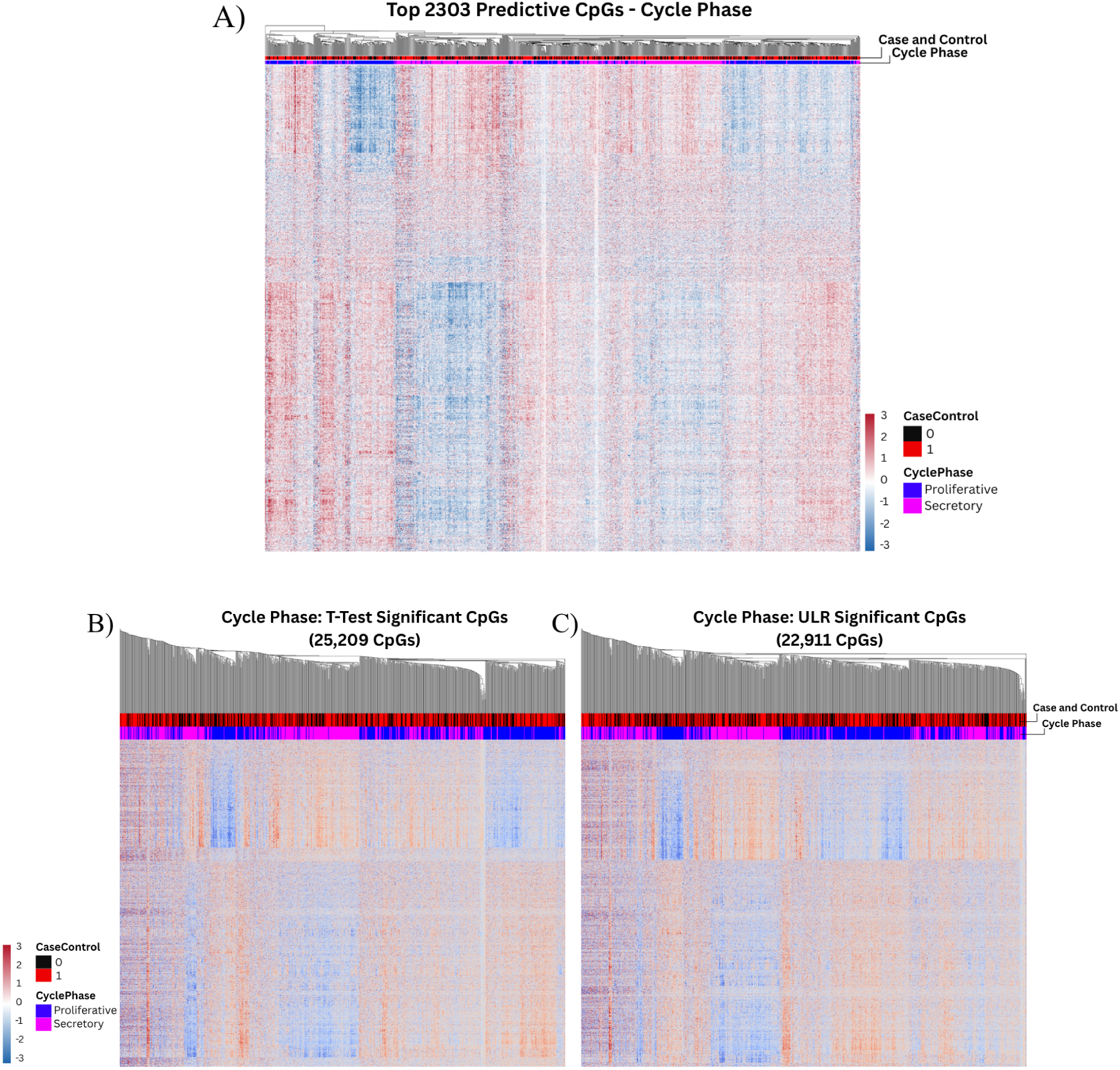
A) Heatmaps of the top predictive CpG sites identified through ridge regression coefficient–based feature selection. Heatmap displays standardized methylation values (row z-scores) for the top 2,303 CpG sites selected from the menstrual cycle phase model. Columns represent individual samples and rows represent CpG sites. Sample annotations indicate menstrual cycle phase (proliferative/secretory). The cycle phase heatmap exhibits coordinated methylation patterns that clearly separate proliferative and secretory phase samples, reflecting the strong influence of hormonal cycling on the endometrial methylome. B and C show heatmaps of CpG sites identified by univariate feature selection methods. Heatmaps display standardized methylation values (row z-scores) for CpG sites identified as significant by false discovery rate (FDR)-corrected univariate analyses. Panel B shows CpGs associated with menstrual cycle phase identified using t-tests (25,209 significant CpGs) and panel C shows the CpGs associated with menstrual cycle phase identified using univariate logistic regression (22,911 significant CpGs). Columns represent individual samples and rows represent CpG sites, with sample annotations indicating endometriosis status and menstrual cycle phase. The cycle phase heatmaps reveal extensive, coordinated methylation patterns that clearly distinguish proliferative and secretory phase samples.

Pathway enrichment analyses identified extensive biological enrichment among cycle phase-associated CpGs. T-test-derived CpGs produced 222 significant Gene Ontology pathways, 13 KEGG pathways, and 4 Reactome pathways following FDR correction. Univariate logistic regression-derived CpGs produced 164 significant Gene Ontology pathways and 16 significant KEGG pathways. Enriched pathways were associated with cellular proliferation, tissue remodeling, developmental regulation, signal transduction, and other processes involved in endometrial function throughout the menstrual cycle (Figure 4). The large number of significant pathways, combined with strong classification performance and extensive differential methylation, indicates that menstrual cycle phase is associated with widespread methylation differences across the endometrium.

**Figure 4.**
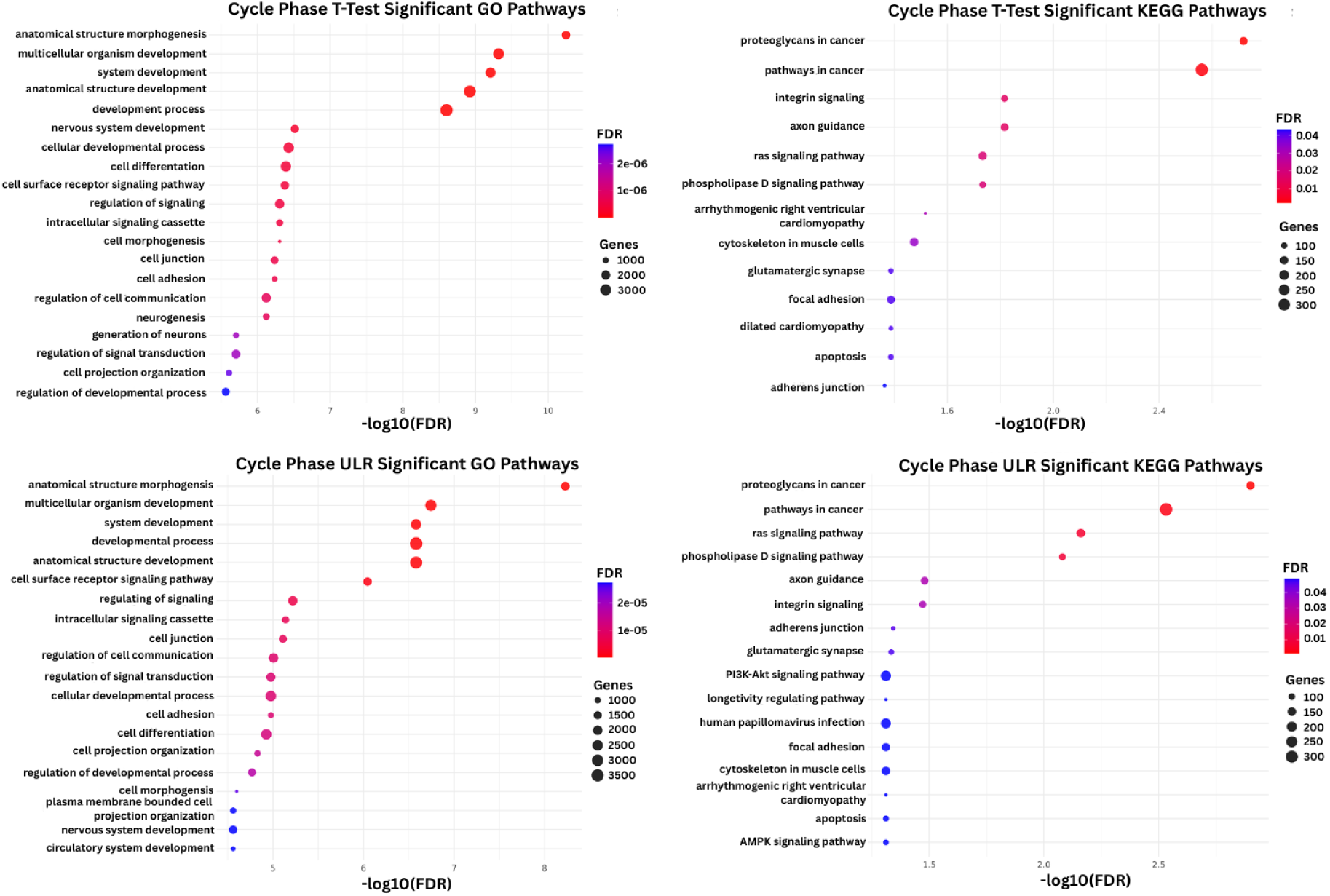
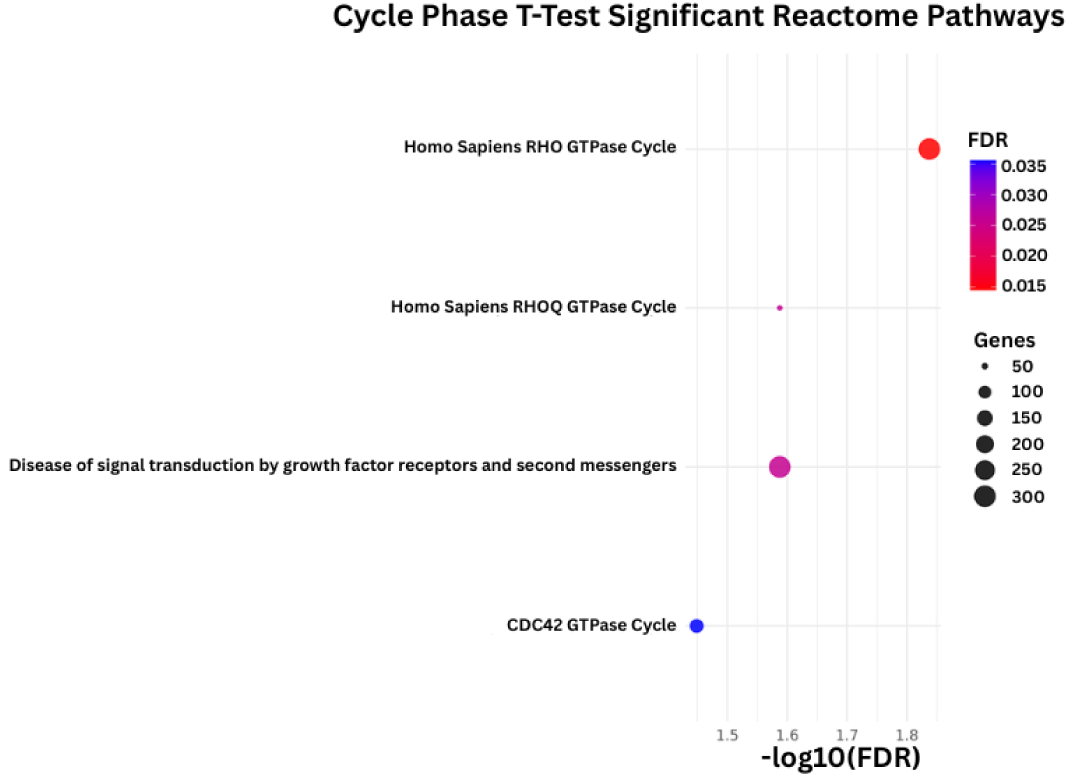
Pathway enrichment analyses for menstrual cycle phase-associated CpG feature sets. Panels A–C show GO, KEGG, and Reactome pathway enrichment results generated using statistically significant CpGs identified through t-test analyses. Panels D and E show GO and KEGG pathway enrichment results generated using statistically significant CpGs identified through univariate logistic regression analyses.

### Machine Learning Reveals a Distributed Predictive Endometriosis Methylation Signature Despite Limited Univariate Associations

The full-feature ridge logistic regression model trained using all 759,345 CpGs classified endometriosis cases and controls with moderate predictive performance. The model achieved a mean cross-validation AUROC of 0.854 and a mean cross-validation AUPRC of 0.903 within the training data. When evaluated on the independently held-out test set, it achieved a held-out test AUROC of 0.875 and a held-out test AUPRC of 0.930 (Figure 5A). Performance remained substantially above random classification and was consistently observed across validation folds. Shuffled-label analyses produced AUROC values near 0.5, indicating that comparable predictive performance was not observed after disruption of the relationship between methylation profiles and outcome labels (Supplementary Figure S2). Although classification performance was lower than that observed for menstrual cycle phase, the model retained substantial predictive ability despite the absence of strong univariate signals. These findings suggest that disease-associated information is present within the methylation data but is not concentrated within a small number of highly significant CpGs.

**Figure 5.**
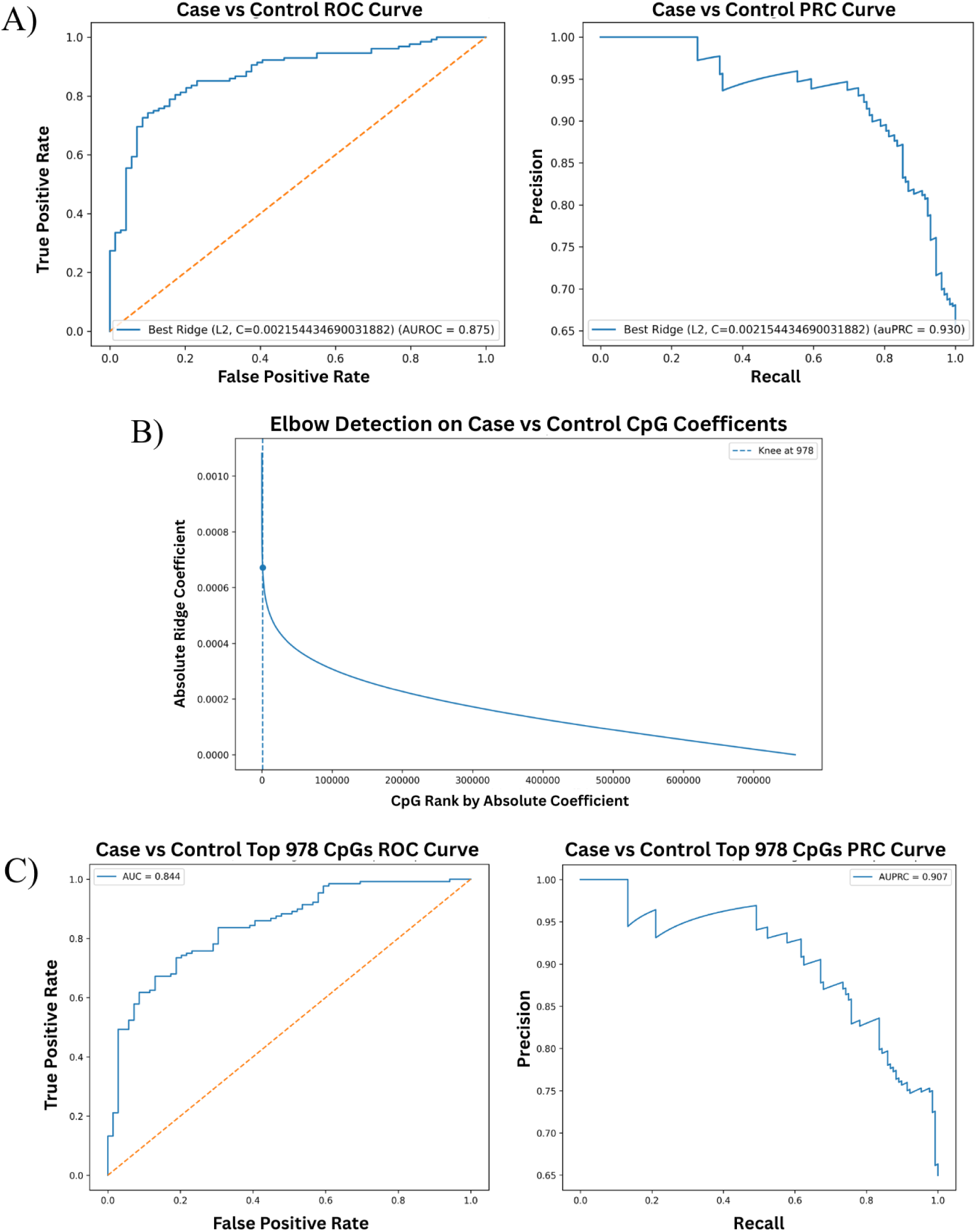
A) Predictive performance of ridge logistic regression models using the full genome-wide DNA methylation feature set. ROC and PRC curves are shown for classification of endometriosis case-control status using all available CpG sites following quality control and SmartSVA correction. The case-control model achieved an AUROC of 0.875 and an AUPRC of 0.930, demonstrating good discrimination between endometriosis cases and controls. B) Elbow point identification for ridge regression coefficient–based feature selection. Absolute ridge regression coefficients were ranked from largest to smallest for the endometriosis case-control model using the full genome-wide methylation feature set. The elbow point was identified using the KneeLocator algorithm to identify where ridge coefficients began to decrease more gradually. The detected elbow corresponded to 978 CpG sites for the case-control model, indicated by the dashed vertical line. This CpG subset was selected as a reduced feature set for downstream modeling and biological interpretation. C) Performance of reduced-feature ridge regression models using CpG subsets selected by coefficient-based elbow analysis. ROC and PRC curves are shown for ridge logistic regression models trained using the top 978 CpG sites identified from the endometriosis case-control model. Following dimensionality reduction from the original ∼759,000 CpG sites, the case-control model achieved an AUROC of 0.844 and an AUPRC of 0.907, indicating that classification performance was largely retained despite the substantial reduction in feature space.

Ranking CpGs by absolute ridge coefficient magnitude identified a reduced feature set consisting of 978 CpGs following elbow detection (Figure 5B). In contrast, independent univariate analyses identified very few statistically significant loci, with per-CpG t-test analyses identifying only five CpGs passing the false discovery rate significance threshold and univariate logistic regression identifying only two significant CpGs. This pattern differed substantially from the menstrual cycle phase analyses, where tens of thousands of CpGs reached statistical significance. The marked discrepancy between multivariate predictive performance and univariate significance suggests that endometriosis-associated methylation differences are distributed across many loci with individually modest effects rather than driven by a small number of strongly associated CpGs. These findings support the use of multivariate approaches that aggregate information across correlated CpGs to capture disease-associated methylation patterns.

A ridge logistic regression model trained using the 978 ridge-selected CpGs retained most of the predictive performance observed with the full-feature model. Following dimensionality reduction from approximately 759,000 CpGs to 978 CpGs, the reduced-feature model achieved an AUROC of 0.844 and an AUPRC of 0.907 (Figure 5C). These findings indicate that much of the disease-associated predictive information was retained despite substantially reducing the feature space.

Heatmaps generated from ridge-selected, t-test-selected, and univariate logistic regression-selected CpGs demonstrated substantially weaker clustering by disease status than was observed for menstrual cycle phase (Figure 6). Samples did not separate as clearly according to endometriosis status, regardless of feature selection method. Similar diffuse methylation patterns were observed across the heatmaps despite large differences in feature count between the ridge-selected and univariate feature sets. The absence of strong disease-associated clustering suggests that endometriosis-related methylation differences are distributed across many loci with individually modest effects rather than concentrated within a small number of highly altered CpGs.

**Figure 6.**
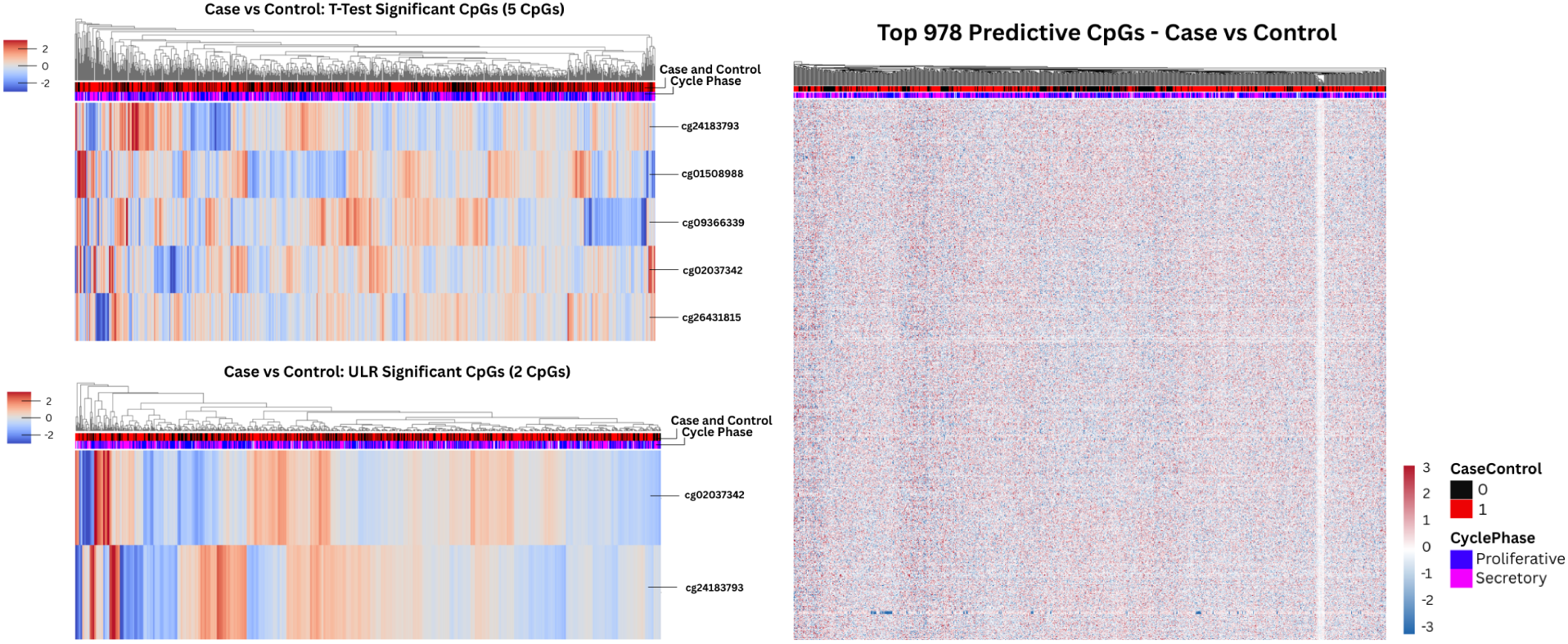
Heatmaps of CpG sites identified by univariate feature selection methods. Heatmaps display standardized methylation values (row z-scores) for CpG sites identified as significant by false discovery rate (FDR)-corrected univariate analyses. Panel A shows CpGs associated with endometriosis case-control status identified using univariate logistic regression (2 significant CpGs) and t-tests (5 significant CpGs). Columns represent individual samples and rows represent CpG sites, with sample annotations indicating endometriosis status and menstrual cycle phase. The heatmap in Panel B shows the top predictive CpG sites identified through ridge regression coefficient–based feature selection. It displays standardized methylation values (row z-scores) for the top 978 CpG sites selected from the endometriosis case-control model. Columns represent individual samples and rows represent CpG sites. Sample annotations indicate endometriosis status (case/control). The case-control heatmap shows subtle clustering across samples.

Pathway enrichment analysis of the case-control CpG sets identified no significantly enriched Gene Ontology, KEGG, or Reactome pathways following false discovery rate correction. The absence of pathway-level enrichment was consistent across both the per-CpG t-test- and univariate logistic regression-derived CpG sets. Pathway enrichment analysis of the 978 ridge-selected CpGs similarly identified no significant GO or KEGG pathways following FDR correction, further supporting the diffuse nature of endometriosis-associated methylation differences. Together with the weak clustering observed in the heatmaps, these findings suggest that disease-associated methylation differences are distributed across many CpGs with individually modest effects rather than concentrated within a few highly altered loci or biological pathways (Figure 6).

## Discussion

The primary finding of this study was that menstrual cycle phase represents the dominant source of methylation variation within eutopic endometrial tissue. Across all analyses, cycle phase produced stronger and more consistent signals than endometriosis status, highlighting the substantial influence of normal endometrial biology on genome-wide methylation patterns. The impact of menstrual cycle phase was evident regardless of the analytical approach used. Samples consistently separated according to proliferative and secretory phase, thousands of CpGs were identified through univariate analyses, and pathway enrichment analyses revealed extensive biological differences between cycle phases. These findings are consistent with the dynamic hormonal regulation and tissue remodeling that occur throughout the menstrual cycle and reinforce the importance of accounting for cycle phase when studying endometrial methylation.

In contrast, the methylation signal associated with endometriosis was much more subtle. Only a small number of CpGs reached significance in the univariate analyses, and pathway enrichment analyses did not identify robust disease-associated pathways. Despite this, ridge regression was still able to distinguish cases from controls with good predictive performance. This discrepancy suggests that disease-associated information is not concentrated within a few highly altered CpGs, but instead is distributed across many loci with individually small effects. Together, these findings suggest that endometriosis is associated with a diffuse and heterogeneous epigenetic signature rather than a small set of large-effect methylation changes.

Our findings are consistent with previous studies of endometrial methylation in endometriosis. Saare et al. reported that samples clustered primarily according to menstrual cycle phase rather than disease status and identified relatively few disease-associated methylation differences. Houshdaran et al. similarly found that many of the strongest methylation changes in eutopic endometrium were phase-dependent. Rahmioglu et al. demonstrated that menstrual cycle phase contributes substantially to variation in endometrial methylation profiles and suggested that much larger sample sizes may be required to reliably detect disease-associated effects.

Perhaps most notably, Mortlock et al., whose dataset formed the foundation of this analysis, also found that cycle phase explained considerably more methylation variation than endometriosis status and identified very few significant disease-associated CpGs. The results of the present study closely mirror these observations while extending them through the use of machine learning approaches, showing that disease-associated patterns remain detectable using multivariate models even when few individual CpGs reach statistical significance. Several aspects of this study strengthen these conclusions. The analysis leveraged one of the largest endometrial methylation cohorts examined to date, incorporated samples from multiple institutions, and evaluated multiple feature selection strategies. SmartSVA correction reduced clustering associated with technical variables while retaining cycle phase-associated variation, and the use of a leakage-safe modeling framework helped ensure that performance estimates remained realistic and reproducible.

Pathway enrichment analysis of menstrual cycle phase also showed strong concordance with prior studies. Shared pathways with Mortlock et al. included focal adhesion, adherens junction, PI3K-Akt signaling, Ras signaling, phospholipase D signaling, apoptosis, axon guidance, and the Reactome RHO GTPase Cycle and CDC42 GTPase Cycle, all of which are involved in cell adhesion, cytoskeletal organization, intracellular signaling, and tissue remodeling during the transition from the proliferative to secretory phase. These findings are further supported by transcriptomic studies identifying focal adhesion, PI3K-Akt signaling, and cell adhesion molecule pathways as key features distinguishing proliferative and secretory endometrium (Yu et al., 2021), as well as enrichment of Gene Ontology terms related to morphogenesis, development, cell differentiation, and tissue remodeling (Apostolov et al., 2024; Mortlock et al., 2022). Together, these overlaps show that the pathways identified here are consistent with previous methylation and transcriptomic studies of the endometrium.

This study also has several limitations. SmartSVA was applied separately to the training and held-out test sets to prevent information from the test samples from influencing model development. Although this approach may result in slightly different latent technical factors being estimated across the two subsets, it prioritizes prevention of information leakage.

Endometriosis is a highly heterogeneous disease, and important biological differences may be obscured when all cases are analyzed together. Although batch correction reduces technical variation, some residual heterogeneity likely remains. External validation was not performed because comparably large, independent endometrial DNA methylation datasets with well-characterized endometriosis status and menstrual cycle phase annotations are limited. Future validation in an independent cohort will therefore be important for evaluating the generalizability of these findings. Cell-type composition was not directly accounted for in this analysis; therefore, some of the observed methylation differences, particularly across menstrual cycle phases, may reflect differences in the types and proportions of cells present in the tissue. Although these findings may have implications for the development of methylation-based biomarkers for endometriosis, the strong influence of menstrual cycle phase on endometrial methylation highlights the importance of accounting for cycle phase when identifying disease-associated methylation patterns (Saare et al., 2016). Further validation in independent cohorts will be needed to determine whether these patterns have potential clinical utility.

Overall, these results suggest that normal physiological changes across the menstrual cycle exert a far greater influence on the endometrial methylome than endometriosis status. At the same time, the ability of multivariate models to distinguish cases from controls indicates that disease-associated methylation differences are present, but are distributed across many CpGs with individually modest effects. Future studies incorporating additional molecular data, external validation cohorts, and more refined patient stratification may help further clarify the biological basis of these diffuse disease-associated methylation patterns.

## Conclusion

We used genome-wide DNA methylation data from 984 eutopic endometrial samples to investigate epigenetic differences associated with menstrual cycle phase and endometriosis.

Across all analyses, menstrual cycle phase emerged as the strongest driver of methylation variation, producing widespread differences across the methylome, extensive pathway enrichment, and highly accurate classification of proliferative and secretory endometrium. These findings highlight the dynamic nature of the endometrium and reinforce the importance of accounting for cycle phase when studying endometrial biology and disease. Although endometriosis-associated methylation differences were far less pronounced, they remained detectable through multivariate modeling. While only a small number of individual CpGs reached statistical significance, ridge regression models were able to distinguish cases from controls with good predictive performance. Together, these findings suggest that endometriosis is associated with a diffuse epigenetic signature distributed across many CpGs rather than a small set of methylation changes.

## Data availability

The DNA methylation data analyzed in this study were previously generated and described by Mortlock et al. (2023) and are publicly available through the NCBI Gene Expression Omnibus under accession GSE223817. The processed dataset used as the starting point for the present analyses was derived from these data as described in the Methods.

## Acknowledgements

We gratefully acknowledge all the women who participated in our studies for their invaluable contribution of biological samples and clinical data, without which this research would not have been possible. We extend our thanks to our clinical and surgical collaborators, particularly Camran Nezhat, Jessica Opoku-Anane, Jeannette Lager, and Alison Jacoby, and consultant gynaecological surgeons Kirana Arambage, Martin Hirsch, Prasanna Supramaniam and Pedro Melo, alongside the trainees and fellows who contributed their clinical expertise and support. We further recognise the efforts of the research team, including Kurtis Garbutt, Kelly Barrett, Lisa Buck, Gavin Collett, Sarah Hutchinson, Sarah Myers, Emily-Jane Philips, Alison Vitonis, and Kim Chi Vo for their commitment to the successful delivery of our studies.

## Author contributions

A.N. performed the computational analyses, interpreted the results, generated figures, and wrote the original draft of this manuscript. U.K. provided mentorship and assisted with review and validation of the analytical approach. C.S., K.L., and M.S. served on A.N.’s thesis committee and provided guidance on the analytical approach and interpretation of results throughout the study. M.S. and L.G. supervised the work. P.G., A.S., I.K., S.Mo., S.H., N.R., S.Mi., K.Z., G.M., C.B., P.R., and J.I. contributed as part of the original analysis team to the generation, curation, analysis, or interpretation of the source dataset used in this study. T.O. co-supervised the present work and provided feedback on the analysis and manuscript. All authors reviewed and revised the manuscript and approved the final version.

