## Supplementary Figures and Captions for "Predicting Endometriosis Status and Menstrual Cycle Phase Using DNA Methylation"

**Supplementary Information**

| Plot Label | Pre | Post | Change |
|---|---|---|---|
| Institute for Analysis | 0.0182 | -0.0895 | -0.1076 |
| Batch | 0.0505 | -0.0225 | -0.0729 |
| Endometriosis (Yes/No) | 0.0028 | 0.0135 | 0.0107 |
| Cycle phase for Analysis | -0.0275 | -0.0893 | -0.0618 |

Supplementary Table S1. Silhouette scores before and after SmartSVA correction for technical (institute of analysis and batch) and biological (endometriosis status and menstrual cycle phase) variables. Following correction, silhouette scores for institute of analysis decreased from 0.0182 to -0.0895 and batch decreased from 0.0505 to -0.0225, indicating reduced clustering by technical variables. The silhouette score for endometriosis status remained near zero (0.0028 to 0.0135), consistent with weak overall clustering by disease status. The silhouette score for menstrual cycle phase changed from -0.0275 to -0.0893.

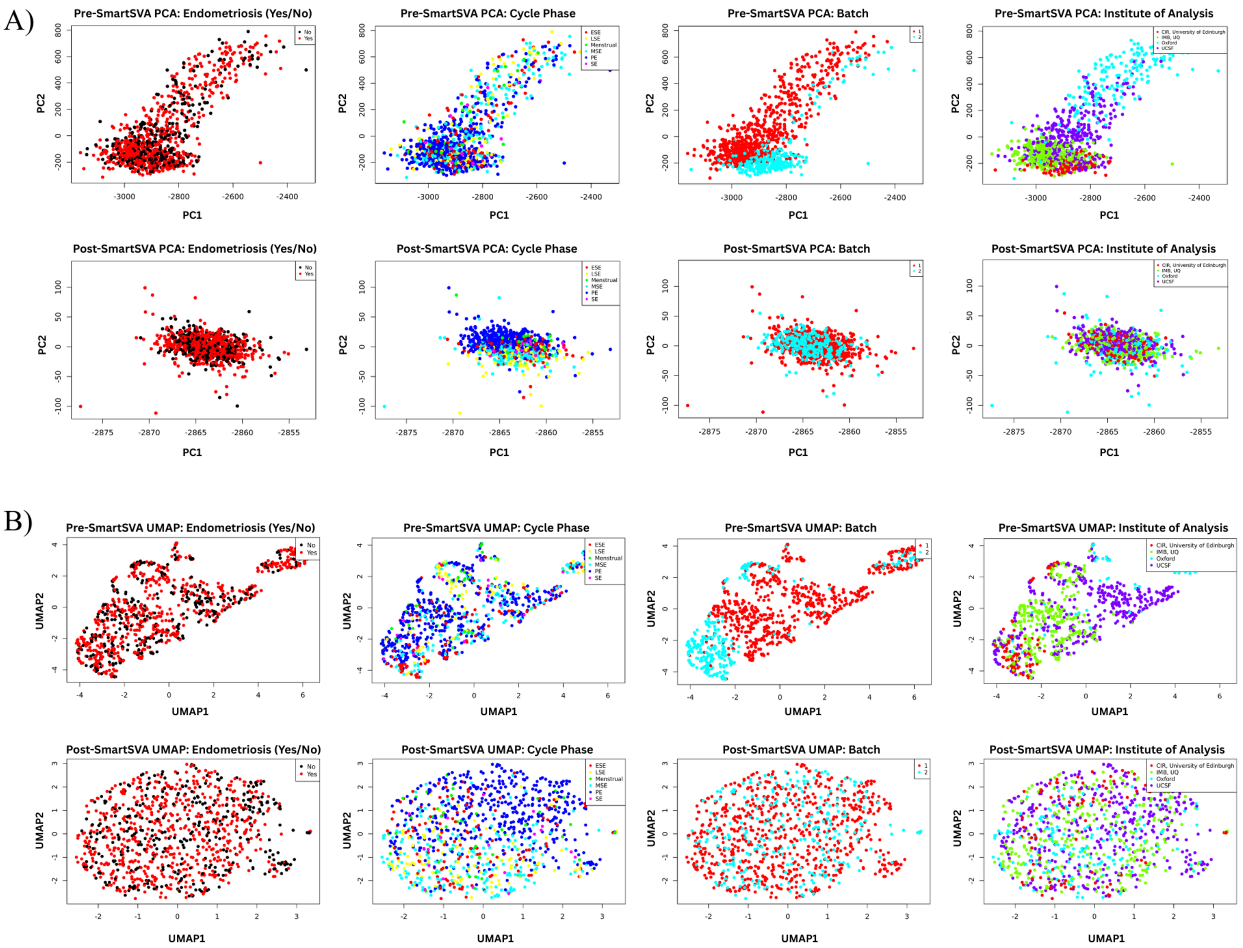


Supplementary Figure S1. PCA and UMAP visualizations before and after SmartSVA correction demonstrating reduction of technical clustering while preserving biological variation. Panel A shows PCA plots colored by endometriosis status, menstrual cycle phase, institute of analysis, and batch before and after SmartSVA correction. Panel B shows UMAP projections colored by institute of analysis, batch, endometriosis status, and menstrual cycle phase before and after SmartSVA correction. SmartSVA substantially reduced clustering associated with batch and institute of analysis while preserving biological variation associated with menstrual cycle phase and, to a lesser extent, endometriosis status.

A)

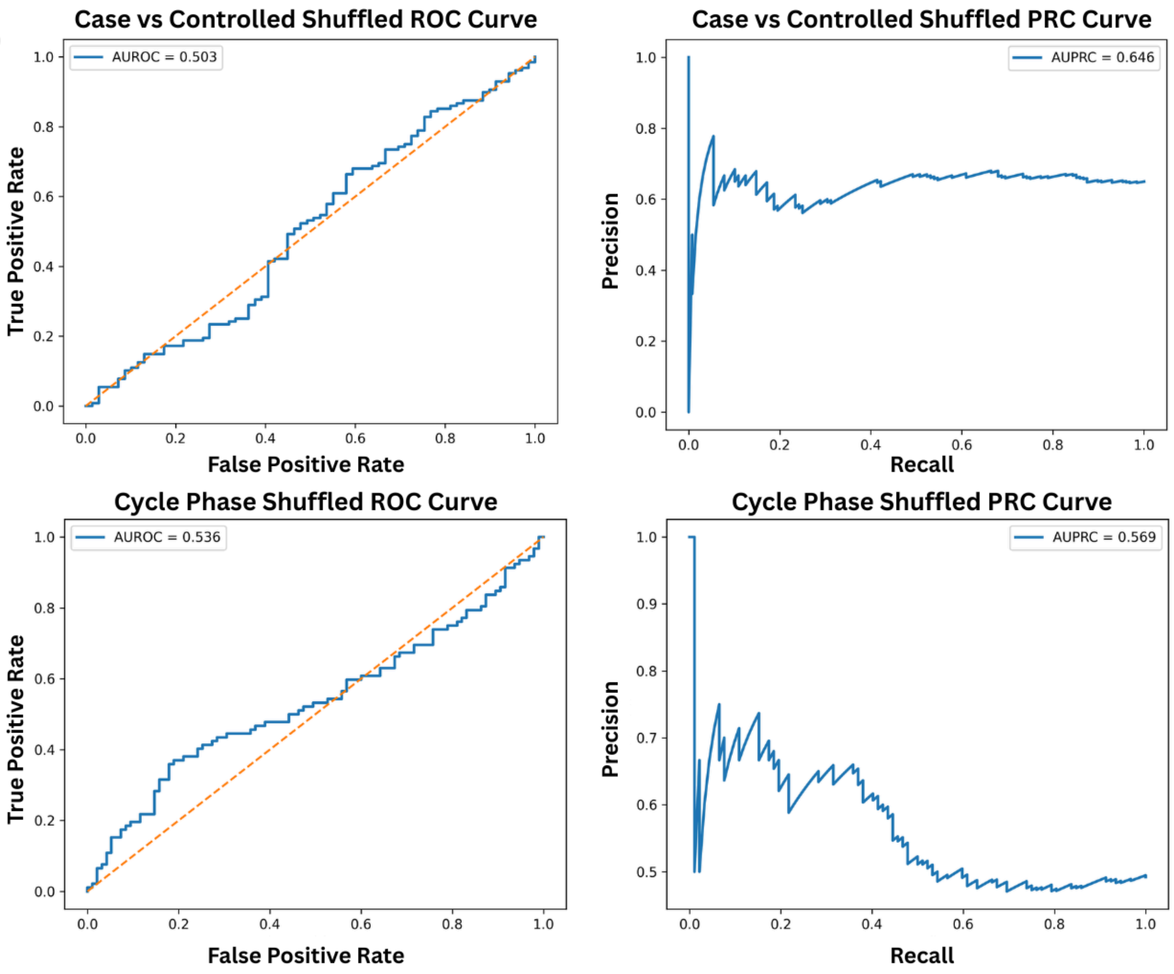


Supplementary Figure S2. ROC and PRC curves for ridge logistic regression models trained using randomly shuffled labels for endometriosis status and menstrual cycle phase. Models were trained using the full feature set following the shuffled-label procedures described in the Methods. As expected, shuffled-label models demonstrated near-random performance (case-control: AUROC = 0.503, AUPRC = 0.646; cycle phase: AUROC = 0.536, AUPRC = 0.569), indicating that comparable predictive performance was not observed after randomization of the outcome labels.
